# Ile35 residue of SOD1 plays a key role in aggregate formation and eye degeneration in a *Drosophila* model of ALS

**DOI:** 10.64898/2026.09.13.751333

**Authors:** Kyoka Yano, Yuzo Fujino, Yoshitaka Nagai, Noriko Fujiwara

**Affiliations:** Department of Biochemistry, School of Medicine, Hyogo Medical University, Nishinomiya, Hyogo 663-8501, Japan; Department of Neurology, Kindai University Faculty of Medicine, Sakai, Osaka 590-0197, Japan; Life Science Research Institute, Kindai University, Sakai, Osaka 590-0197, Japan

**Keywords:** Amyotrophic lateral sclerosis (ALS), Cu/Zn-superoxide dismutase (SOD1), *Drosophila*, Aggregate

## Abstract

- The pathophysiology of ALS caused by mutant SOD1 remains unclear.
- We previously demonstrated, using the BioID2-EGFP sandwich expression system in cultured cells, that the Ile35 residue of ALS-linked mutant SOD1 promotes its aggregate formation.
- However, the precise role of the SOD1 Ile35 residue in aggregate formation and neurodegeneration *in vivo* remained unclear.
- *Drosophila* expressing BioID2-G93A-EGFP demonstrated aggregate formation and eye degeneration with reduced eye size, whereas those expressing BioID2-G93A/I35S-EGFP showed less of these phenotypes.
- These findings highlight the crucial role of the Ile35 residue of ALS-linked SOD1 in aggregate formation and neurodegeneration *in vivo*.

---

Amyotrophic lateral sclerosis (ALS) is a fatal neurodegenerative disease characterized by the degeneration of upper and lower motor neurons, leading to progressive motor dysfunction and respiratory muscle weakness [1]. Although only 10% of ALS patients have a family history of the disease, more than 40 ALS-associated genes have been identified to date by large-scale genetic studies [2], since the first causative gene for ALS (Cu/Zn-superoxide dismutase (SOD1)) was discovered in 1993 [3]. SOD1 is a homodimeric antioxidant enzyme composed of 153 amino acids and binds to one copper ion and one zinc ion per subunit. To date, more than 220 mutations in the *SOD1* gene have been identified as causing ALS (http://alsod.iop.kcl.ac.uk). However, despite extensive research, the mechanisms responsible for the motor neuron degeneration caused by ALS-linked SOD1 mutations have remained unclear to date.

The misfolding of proteins and the formation of inclusions containing protein aggregates are major pathological features of neurodegenerative diseases, such as Alzheimer’s disease, Parkinson’s disease, and ALS [4]. ALS mutant SOD1 proteins are more prone to misfolding and dimer destabilization, and more readily form insoluble aggregates [5]. We previously reported that the Ile35 residue in the hydrophobic region of SOD1 plays a crucial role in the intracellular aggregate formation of ALS-linked SOD1, using the BioID2-EGFP sandwich expression system. In fact, substituting Ile35 with serine (I35S) significantly decreased the extent of aggregation of the A4V, H46R, and G93A mutant SOD1 as well as wild type (WT) SOD1 [6]. McAlary et al. also reported that replacing Ile35 with proline (I35P) decreased the aggregation propensity and toxicity of G85R mutant SOD1 [7]. However, as these findings are based on the aggregate formation observed in cultured cells, the role of the Ile35 residue of SOD1 in aggregate formation and neurodegeneration *in vivo* remains unclear.

*Drosophila melanogaster* (the common fruit fly) has a short life cycle and produces many offspring; therefore, it is a powerful model for analyzing the molecular basis of neurodegenerative diseases, including the polyglutamine diseases, Parkinson’s disease and ALS [8]. Transgenic flies expressing ALS-linked mutant SOD1 recapitulate many features of ALS pathology: SOD1 aggregation, mitochondrial dysfunction, locomotor dysfunction, and reduced survival [9]. ALS model flies expressing other ALS-associated proteins such as Fused in Sarcoma (FUS) and TAR DNA-binding protein 43 (TDP43) have also been reported to show protein aggregation and neurodegenerative phenotypes including locomotor dysfunction, reduced survival, and compound eye degeneration [10].

In this study, we conducted a comparative analysis of *Drosophila* ALS models expressing BioID2-SOD1 (G93A)-EGFP (high-aggregate-forming type) and BioID2-G93A/I35S-EGFP (low-aggregate-forming type) to elucidate the role of the Ile35 residue of SOD1 in aggregate formation and neurodegeneration, as well as the association between SOD1 aggregate formation and neurodegeneration *in vivo*.

Transgenic fly lines carrying the following transgenes were generated: *UAS-EGFP* (hereinafter EGFP), *UAS-BioID2-EGFP* (hereinafter Bio-EGFP), *UAS-BioID2-SOD1 (WT)-EGFP* (hereinafter Bio-WT), *UAS-BioID2-SOD1 (G93A)-EGFP* (hereinafter Bio-G93A), and *UAS-BioID2-SOD1 (G93A/I35S)-EGFP* (hereinafter Bio-G93A/I35S). These transgenes were expressed under the control of the eye-specific *GMR-Gal4* driver at 28 °C [11].

To evaluate aggregate formation, eye imaginal discs were dissected from third-instar larvae, fixed with 4% paraformaldehyde, and mounted onto slides. The number of EGFP-positive aggregates in the eye discs was counted using a confocal laser-scanning microscope. Female adult flies were used for the evaluation of eye phenotype. Scanning electron microscopy images of 1 to 2-day-old flies were used to evaluate ocular degeneration, and stereomicroscopy images of 1 to 3-day-old flies were used for quantifying eye size. Detailed experimental procedures are described in the Supplemental Methods.

We first confirmed the expression of the proteins encoded by the constructed transgenes at comparable levels and their expected molecular weights in the heads of *Drosophila* by immunoblotting (Fig. S1). Bio-WT, Bio-G93A, and Bio-G93A/I35S proteins demonstrated a band corresponding to BioID2-SOD1-EGFP (molecular weight: 70 kDa) and reacted with an anti-SOD1 antibody. We next analyzed aggregate formation in the eye imaginal discs of larvae expressing these transgenes. As shown in Figures 1A and 1B, numerous aggregates, with a mean of 25 aggregates per 15 ommatidia, were observed in Bio-G93A flies (arrows in the enlarged view), predominantly in the cytoplasm. In contrast, Bio-WT and Bio-G93A/I35S flies showed diffuse EGFP fluorescence predominantly in the nucleus, with only a small number of cytoplasmic aggregates (Fig. 1A). These findings suggest that Bio-G93A proteins translocate from the nucleus to the cytoplasm and form aggregates, and that the additional I35S mutation suppressed the aggregate formation of Bio-G93A. This result is consistent with our previous findings from the expression of these genes in cultured cells [6]. Misfolded SOD1, such as the G93A mutant, tends to be exported from the nucleus via a nuclear export signal-like sequence that is normally buried within the SOD1 protein [12].

**Figure 1.**
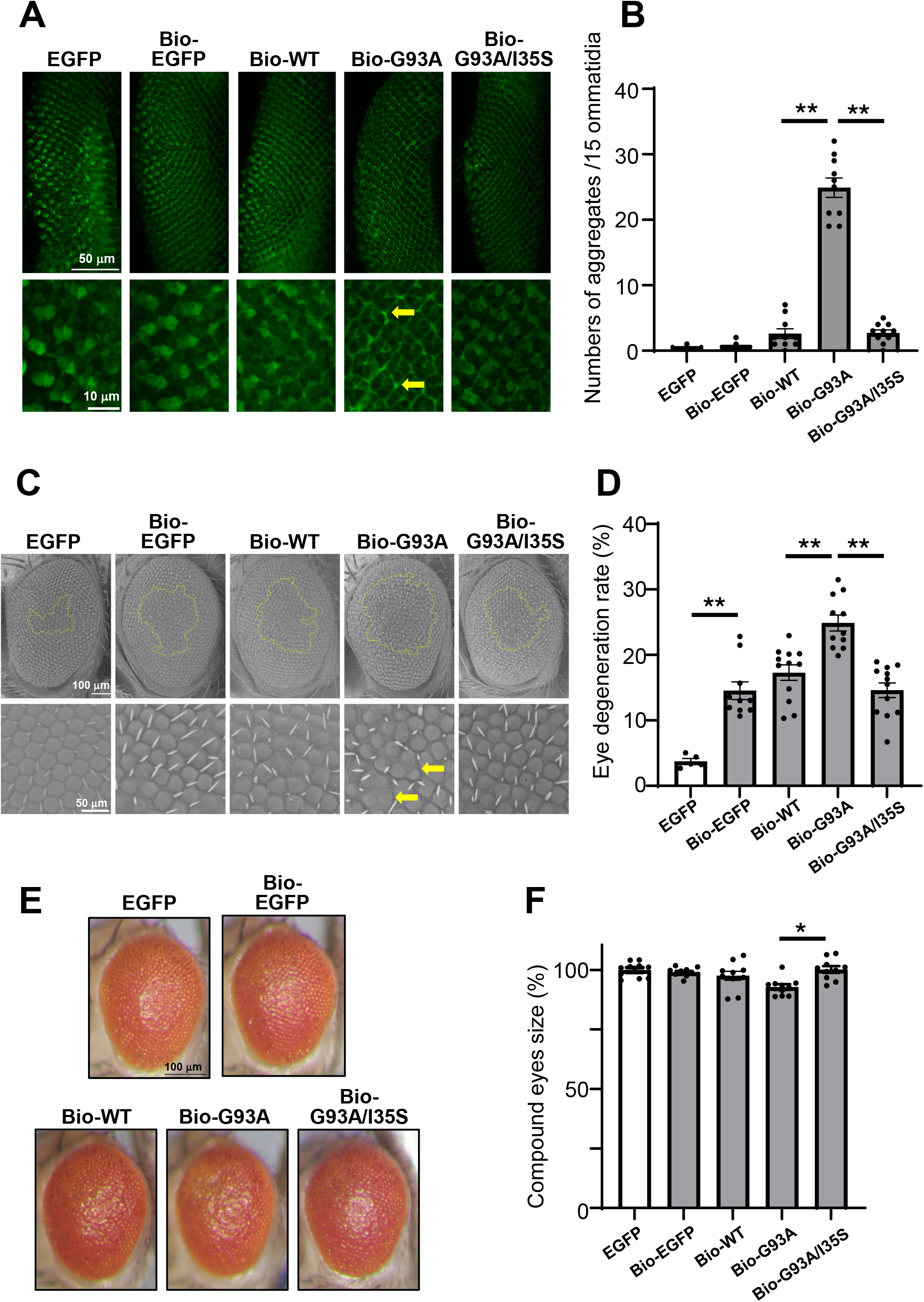
BioID2-G93A-EGFP induces aggregate formation and eye degeneration in a *Drosophila* ALS model, whereas the additional I35S mutation suppresses these effects. **(A)** Aggregates formed in the eye imaginal discs of fly larvae. Each lower panel shows a higher magnification image of the corresponding upper panel. **(B)** The number of aggregates in 15 ommatidia were counted for each fly line. Data are presented as the mean ± SEM. (n = 10) Statistical analyses were performed using one-way ANOVA followed by Tukey’s post-hoc test. \*\**p* < 0.001 **(C)** Fused eye morphology of the adult compound eye (aged 1–2 days) observed using a scanning electron microscope. The fused eye areas are marked with yellow lines in the upper panel. Each lower panel shows a higher magnification image of the corresponding upper panel. **(D)** Rates of eye fusion (%) were expressed as the ratio of the fused eye area to the total area of the compound eye. Data are presented as the mean ± SEM. (n = 5 to 12) Statistical analyses were performed using one-way ANOVA followed by Tukey’s post-hoc test. \*\**p* < 0.001 **(E)** Stereomicroscopy images of fly eyes 1 to 3 days post-eclosion. **(F)** Compound eye sizes (%) were expressed as relative values, with the eye size of flies expressing EGFP only set at 100%. Data are presented as the mean ± SEM. (n = 10) Statistical analyses were performed using one-way ANOVA followed by Tukey’s post-hoc test. \**p* < 0.01

We next investigated whether the expression of these proteins induces neurodegeneration *in vivo*, by scanning electron microscopy. In Bio-G93A flies, some ommatidia in the compound eyes were fused, consistent with ocular neurodegeneration (Fig. 1C, arrows in the enlarged view), and the fused area accounted for approximately 25% of the eye surface area (Fig. 1D). By contrast, the fused eye area was smaller in the Bio-G93A/I35S flies and was comparable to Bio-WT flies (Figs. 1C and 1D), suggesting that the additional I35S mutation suppresses the eye degeneration induced by Bio-G93A. Although slight ommatidial fusion was also observed in control EGFP flies, this mild degeneration may reflect the intrinsic toxicity of the *GMR-GAL4/UAS* system at 28 °C [11]. Consistent with these observations, the stereomicroscopy images showed that the eyes of Bio-G93A flies were slightly but significantly smaller than those of the other lines (Figs. 1E and 1F). Eye sizes of the Bio-G93A/I35S flies were restored to levels comparable to those in control EGFP, Bio-EGFP, and Bio-WT flies, which indicates that the additional I35S mutation ameliorates the eye-size abnormality caused by the G93A mutation. Taken together, these findings indicate that eye degeneration is associated with aggregate formation and that the Ile35 residue plays a crucial role in SOD1 aggregate formation and neurodegeneration in these *Drosophila* models.

The Ile35 residue is located in β-strand III and its hydrophobic side chain points toward the interior of the SOD1 molecule and is surrounded by nine hydrophobic amino acid residues within a distance of 3 Å [6]. Recently, two families harboring the I35F mutation have been reported as having a novel fALS mutation [13]. Like isoleucine, phenylalanine is a hydrophobic amino acid, and we have confirmed that BioID2-I35F-EGFP forms aggregates in cultured cells (data not shown), similarly to other fALS variants harboring Ile35 [6]. We therefore consider that the hydrophobicity of Ile35 in SOD1 contributes to its aggregation in organisms, and that the substitution of Ile35 with Ser (I35S) decreases the hydrophobic effect and prevents aggregation and neurodegeneration. In addition, our atomistic molecular dynamics simulations regarding SOD1 structure suggest that structural changes in the hydrophobic region containing Ile35 contribute to the large fluctuations of the electrostatic loop (loop VII) caused by the G93A mutation, and that the additional I35S mutation mitigates these abnormal fluctuations [Fukushima K and Fujiwara N, unpublished work]. Such fluctuations and destabilization of the electrostatic loop are known to be common characteristics of fALS-associated SOD1 mutations [14]. These findings suggest that the activation of electrostatic loop fluctuations, which are mediated by changes in the hydrophobic region containing Ile35, leads to neurodegeneration by enhancing the intermolecular interactions between SOD1 molecules and promoting aggregate formation.

BioID2 is an improved version of the original biotin ligase used in the “Biotin IDentification (BioID)” method, in which the biotin labels and identifies proteins that interact with a target protein within the cell [15]. This method will enable the identification of proteins in the eyes of *Drosophila* that bind to ALS mutant SOD1 (high-aggregate-forming type) but not to I35S mutant SOD1 (low-aggregate-forming type). This approach is expected to elucidate the molecular link between SOD1 aggregation and neurodegeneration. Insights into these molecular mechanisms should contribute to the discovery and development of therapeutic drugs for ALS.

In conclusion, we demonstrated that the Ile35 residue of SOD1 plays a crucial role not only in its aggregate formation but also in neurodegeneration *in vivo*, and that the neurodegeneration is associated with the formation of SOD1 aggregates.

## Abbreviations

ALS: Amyotrophic lateral sclerosis
fALS: familial ALS
SOD1: Cu/Zn-superoxide dismutase
EGFP: Enhanced green-fluorescent protein
ANOVA: Analysis of variance
SEM: Standard error of the mean

## CRediT authorship contribution statement

Kyoka Yano: Investigation, Data curation, Formal analysis, Visualization. Yuzo Fujino: Investigation, Methodology, Formal analysis, Validation. Yoshitaka Nagai: Writing – review & editing, Supervision, Resources. Noriko Fujiwara: Conceptualization Writing – original draft, Writing – review & editing, Project administration, Supervision, Funding acquisition.

## Funding

This work was supported by JSPS KAKENHI Grant Number JP22K11870 and JP25K14826 for Scientific Research (C) (to NF), Grant Number JP24H00630 for Scientific Research (A), JP21H02840 for Scientific Research (B), and JP20H05927 for Transformative Research Areas (A) (Multifaceted Proteins) (to YN), and by Intramural Research Grant Number 3–9 and 6–9 for Neurological and Psychiatric Disorders from the National Center of Neurology and Psychiatry (to YN), and in part supported by Hyogo Medical University Diversity Grant for Research Promotion,2026

## Data availability

Data will be made available on request.

## Competing interests

The authors declare that they have no competing interests.

## Ethical approval

Not applicable

## Acknowledgements

We wish to thank Dr. H Akiko Popiel for critical reading and English editing of the manuscript.

## Notes

### Competing Interest Statement

The authors have declared no competing interest.

