## Supplemental Information for "Ile35 residue of SOD1 plays a key role in aggregate formation and eye degeneration in a *Drosophila* model of ALS"

**Noriko Fujiwara, Ph.D**

Department of Biochemistry, School of Medicine, Hyogo Medical University

1-1 Mukogawa-cho, Nishinomiya, Hyogo 663-8501, Japan,

### Supplemental Methods

#### Fly stocks

All fly stocks were maintained at 25 °C and crosses were performed at 28 °C on standard cornmeal-yeast-glucose medium. Female flies were used in all experiments. The *GMR-Gal4* driver line has been described previously [1].

#### Generation of transgenic flies

To generate transgenic fly lines carrying *UAS-EGFP, UAS-BioID2-EGFP, UAS-BioID2-SOD1 (wild-type (WT))-EGFP, UAS-BioID2-SOD1 (G93A)-EGFP,* and *UAS-BioID2-SOD1 (G93A/I35S)-EGFP,* pUASTattB vectors containing the respective transgenes were injected into fly embryos of the attP2 strain. Site-specific transgenesis was used to insert each construct into the same genomic locus, minimizing variation in positional effects on transgene expression. These transgenic fly lines were generated by BestGene Inc. (Chino Hills, USA) using standard procedures.

#### Quantification of aggregates in eye imaginal discs

Eye imaginal discs were dissected from female third-instar larvae in ice-cold phosphate-buffered saline (PBS) (−), and then fixed with 4% paraformaldehyde in PBS for 30 min. Following three washes with PBS, nuclei were stained with Hoechst 33342. Samples were mounted in SlowFade Gold antifade reagent (Thermo Fisher Scientific, Waltham, USA) and observed using a confocal laser-scanning microscope (FV3000; Olympus, Tokyo, Japan).

The number of EGFP-positive aggregates in eye imaginal discs was quantified in photoreceptor neurons within 15 developing ommatidia in rows 2 and 3 at the posterior edge of the eye disc, identified by Hoechst 33342 staining. These ommatidia were selected because they are at comparable stages of development and can be readily identified. EGFP-positive aggregates were defined as bright fluorescent punctate signals. Ten eye discs per genotype were analyzed using ImageJ software (version 1.54, National Institutes of Health, USA).

**Imaging and quantification of fly eyes**

Compound eyes (5–12 flies per genotype) from female flies (aged 1–2 days) were analyzed for eye degeneration. Scanning electron microscopy was performed using a TM1000 scanning electron microscope (Hitachi, Tokyo, Japan). Ommatidial fusion was quantified as previously described [2]. Briefly, the area containing fused ommatidia was expressed as a percentage of the total compound eye area. Eye sizes of compound eyes from female flies (aged 1–3 days; 10 flies per genotype) were quantified from images obtained using a stereomicroscope (SZX10; Olympus, Tokyo, Japan) equipped with a digital camera (DP23; Olympus, Tokyo, Japan). The obtained images were analyzed using ImageJ software.

**Immunoblotting**

The heads of 5 flies were homogenized in 75 μL of PBS (−) for 10 minutes, sonicated, and then homogenized together with 25 μL of 4 × SDS sample buffer (250 mM Tris-HCl [pH 6.8], 4% SDS, 2.5% glycerol, 0.1% bromophenol blue, and 10% 2-mercaptoethanol). An 8 μL aliquot of the supernatant obtained by centrifuging the protein samples at 15,000

× g for 20 min at 4 °C was separated by SDS-PAGE on a 5% to 20% gradient gel (ATTO, Tokyo, Japan), and then transferred onto a PVDF membrane (Merck Millipore, Burlington, MA, USA) using a Trans-Blot SD Semi-Dry Transfer Cell (Bio-Rad Laboratories, Hercules, CA, USA). The membrane was incubated for 1 h at room temperature in Tris-buffered saline containing 0.05% Tween 20 (TBS-T; 20 mM Tris-HCl, 150 mM NaCl, 0.05% Tween 20, pH 7.6). containing 5% (w/v) skim milk and then incubated with the appropriate antibodies. The target proteins were detected by a chemiluminescence method using Immobilon® Crescendo Western HRP Substrate (Merck Millipore) using an LAS3000 luminescent image analyzer (Fujifilm, Tokyo, Japan). To perform re-probing, antibodies were removed from the membrane using a commercially available Stripping Solution (Fujifilm).

The antibodies used and their dilutions were as follows:

Rabbit anti-GFP (Santa Cruz Biotechnology), 1:3,000

Goat anti-human SOD1 [3], 1:3000

Rabbit anti-β-actin (GeneTex), 1:5,000

HRP-conjugated anti-rabbit IgG (H+L) (Proteintech), 1:5,000

HRP-conjugated anti-goat IgG (Proteintech), 1:5,000

**Data analysis**

All graphs were generated and statistical analyses were conducted using GraphPad Prism software (version 10.6.0, GraphPad Software, Boston, MA, USA). Numerical values are expressed as the mean ± SEM. The statistical analyses were performed using one-way ANOVA test with the Tukey’s post hoc test.

**Additional file**

Figure S1

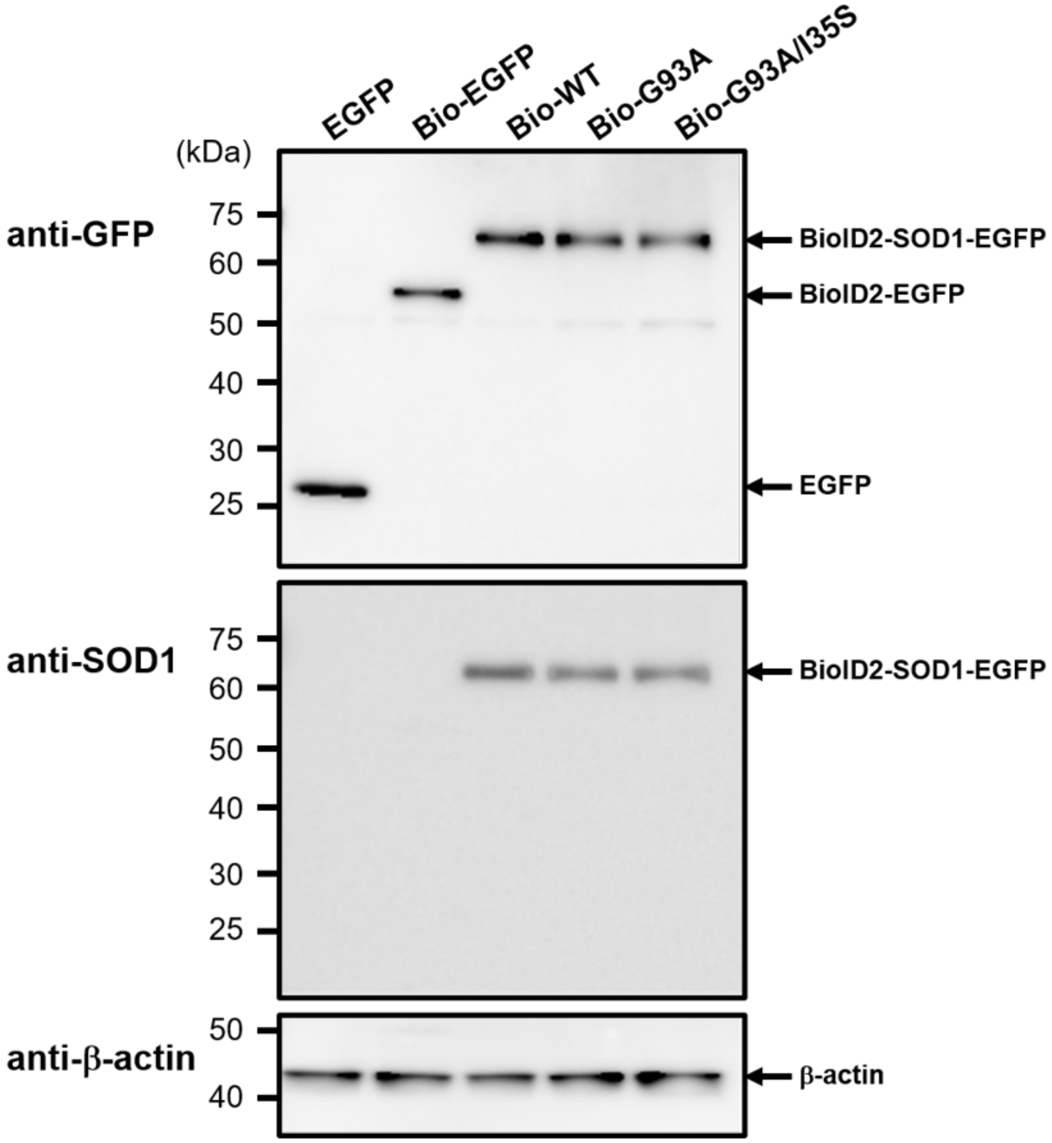


**Figure S1. Expression of BioID2-fused SOD1 proteins from the constructed genes in *Drosophila* heads analyzed by immunoblotting**

The three BioID2-SOD1-EGFP proteins (molecular weight = 70 kDa) reacted with both anti-GFP and anti-SOD1 antibodies. BioID2-EGFP (54 kDa) and EGFP (27 kDa) reacted with only the anti-GFP antibody. β-actin was used as a loading control. The reactions to these antibodies were performed on the same membrane.
